# HNRNPA2B1 controls an unfolded protein response-related prognostic gene signature in prostate cancer

**DOI:** 10.1101/2022.06.21.495112

**Authors:** John G Foster, Esteban Gea, Mosammat A Labiba, Chinedu A Anene, Jacqui Stockley, Celine Philippe, Matteo Cereda, Kevin Rouault-Pierre, Hing Leung, Conrad Bessant, Prabhakar Rajan

**Author notes:** Arquer Diagnostics Ltd, North East Business Innovation Centre, Wearfield, Sunderland, SR5 2TA, UK. Alloy Therapeutics, Inc, 44 Hartwell Ave, Lexington, Massachusetts 02421, USA. **Correspondence to:** Dr. Prabhakar Rajan, Centre for Cancer Cell and Molecular Biology, Barts Cancer Institute, Cancer Research UK Barts Centre, Queen Mary University of London, Charterhouse Square, London, EC1M 6BQ, UK.

## Abstract

HNRNPA2B1 is associated with prostate cancer (PC) disease aggressiveness and underlies pro-tumourigenic cellular stress responses. By analysing >500 PC transcriptomes, we reveal that *HNRNPA2B1* over-expression is associated with poor patient prognosis and stress response pathways. These include the *“protein processing in the endoplasmic reticulum”* (ER) pathway, which incorporates the unfolded protein response (UPR). By RNA-sequencing of HNRNPA2B1-depleted cells PC cells, we identified HNRNPA2B1-mediated down-regulation of UPR genes including the master ER-stress sensor *IRE1*, which induces ER proteostasis. Consistent with *IRE1* down-regulation in HNRNPA2B1-depleted cells, we observed reduced splicing of the IRE1-target and key UPR effector XBP1s. Furthermore, HNRNPA2B1 depletion up-regulates expression of the IRE1-dependent decay (RIDD) target gene *BLOC1S1*, which is degraded by activated IRE1. We identify a HNRNPA2B1-IRE1-XBP1-controlled four gene prognostic biomarker signature (HIX) which classifies a subgroup of primary PC patients at high risk of disease relapse. Pharmacological targeting of IRE1 attenuated HNRNAPA2-driven PC cell growth. Taken together, our data reveal a putative novel mechanism of UPR activation in PC by HNRNPA2B1, which may promote an IRE1-dependent yet potentially-targetable recurrent disease phenotype.

## Introduction

The *HNRNPA2B1* gene codes for two protein isoforms, A2 and B1, which are members of the heterogeneous nuclear ribonuclear protein (HNRNP) family of RNA-binding proteins (RBPs) (Liu & Shi 2021). HNRNPA2B1 modulates cellular phenotypes in disease via multiple different RNA processing functions including alternative pre-mRNA splicing (Li et al 2017) and mRNA stability (Martinez et al 2016). In cancer, HNRNPA2B1 can stabilise (Fahling et al 2006, Stockley et al 2014) or destabilise (Zuccotti et al 2014) mRNAs or control oncogenic splicing switches during tumorigenesis (Clower et al 2010, David et al 2010).

Rapid cellular proliferation during tumorigenesis requires an increased rate of protein synthesis (Lee et al 2021), however a limited oxygen and nutrient supply disrupts proteostasis and causes oxidative stress (Bartoszewska & Collawn 2020). An early cellular response to stress is the stalling of mRNA translation and aggregation of pre-initiation translation complexes into stress granules (Marcelo et al 2021) which recruit RBPs including EWSR1, HNRNPA0, HNRNPA1 and HNRNPA2B1 (Jiang et al 2021, Wolozin & Ivanov 2019). Recent studies have identified HNRNPA2B1 cytoplasmic to nuclear translocation in low oxygen conditions, and its association with the polysome, which contains proteins involved in translation, and regulates proteostasis (Ho et al 2020, Yao et al 2013).

Prolonged stress-induced disruption of cellular proteostasis can lead to increased demand on the protein folding machinery of the endoplasmic reticulum (ER) (Rzymski et al 2010), causing protein re-folding, or destruction of terminally misfolded proteins. ER stress triggers altered unfolded protein response (UPR) gene expression profiles via activation of transcription factor sensors including XBP1, ATF4, and nATF6, which control the three key signalling branches of the UPR (Han & Kaufman 2017). Sustained UPR activation leads to increased tumorgenicity, metastatic potential, and therapy resistance of cancer cells (Cubillos-Ruiz et al 2017). In patients, UPR pathway genes are up-regulated (Han & Kaufman 2017), and the transcriptional targets of XBP1, ATF4 and nATF6 are associated with poor survival (Pallmann et al 2019, Sheng et al 2019).

Prostate cancer (PC) is the commonest male-specific cancer and leading male-specific cause of cancer death (Rebello et al 2021). In PC, proteostasis is disrupted (Bouchard et al 2018), and all three branches of the UPR are activated (Pachikov et al 2021, Pallmann et al 2019, Sheng et al 2019). IRE-1-XBP1 activation leads to initiation of c-MYC dependent transcription and is associated with poor patient prognosis (Sheng et al 2019). In light of evidence implicating HNRNPA2B1 in PC (Stockley et al 2014) and cellular stress (Ho et al 2020, Wolozin & Ivanov 2019, Yao et al 2013), we hypothesised that HNRNPA2B1 may control several stress response pathways including UPR in PC. We reveal for the first time that HNRNPA2B1 regulates expression of UPR pathway genes including *IRE1*, mediates non-canonical splicing of XBP1 mRNA, and controls a gene signature of IRE1-XBP1 activation that is associated with poor PC patient prognosis.

## Results

### HNRNPA2B1 overexpression is associated with poor patient prognosis and cellular stress pathways in prostate cancer

We have previously shown that HNRNPA2B1 protein expression is specifically up-regulated in patients with aggressive prostate cancer (PC) (Stockley et al 2014). To validate these findings, we explored *HNRNPA2B1* expression in RNA sequencing (RNA-Seq) data from primary prostate tumours (n=491) and adjacent benign prostate tissue (n=52) (Sanchez-Vega et al 2018). *HNRNPA2B1* mRNA expression was significantly higher in tumours compared to adjacent benign prostate tissue (Fig. 1A). To determine whether high expression of *HNRNPA2B1* is associated with poor patient prognosis, we stratified tumours into two groups based on the normalized expression levels of *HNRNPA2B1*, with high expression considered the top 25% of the distribution across samples, and the rest of samples considered as low expression. High expression of *HNRNPA2B1* was associated with a statistically significant reduction in patient survival, as compared with patients with low *HNRNPA2B1* expression (Fig. 1B).

**Figure 1.**
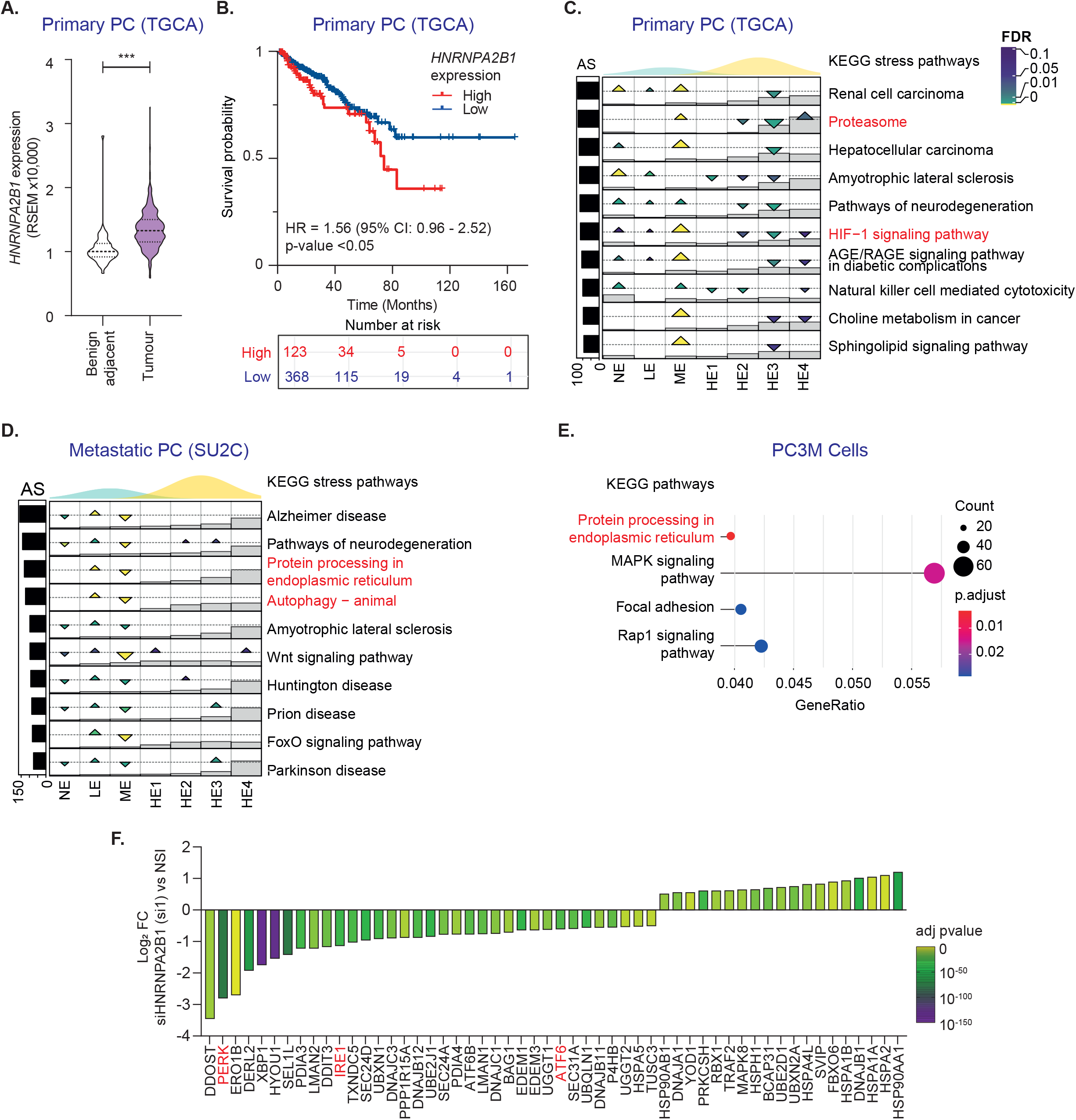
HNRNPA2B1 overexpression is associated with poor patient prognosis and cellular stress pathways in primary prostate cancer. **(A)** Distribution of *HNRNPA2B1* expression values reported as RNA-Seq by Expectation-Maximization (RSEM) in primary prostate tumours and benign adjacent tissue from The Cancer Genome Atlas (TCGA) patient cohort. Two-tailed T-test was used to compare treatment groups. *** = p<0.001 **(B)** Kaplan-Meier plot of disease-free survival for primary PC patients stratified by *HNRNPA2B1* expression (low = <1^st^-3^rd^ quartile and high = >3^rd^ quartile). The number of patients at risk for each group are presented in the table below each X-axis time point. Univariable Cox PH-derived hazard ratios (HR) with 95% confidence intervals (CI) and two-tailed log-rank test p-values are shown. **(C-D)** GSECA analysis performed on **(C)** primary PC (TCGA) and **(D)** metastatic PC (SU2C) RNA-Seq datasets by stratification of cohorts based on *HNRNPA2B1* expression. Genes in a given Kyoto Encyclopaedia of Genes and Genomes (KEGG) pathway are separated into seven expression classes: NE = not expressed, LE= lowly expressed, ME = medium expression, HE1-4 = high expression. Triangles compare the difference in the cumulative proportion of genes in an expression class between *HNRNPA2B1* high and low expression groups, and represent the size and enrichment (up) or depletion (down) of genes. AS = association score. **(E)** KEGG pathway gene enrichment analysis of differentially expressed genes (p<0.05 and absolute log_2_ fold change > 0.5 or < −0.5) identified by RNA-Seq of PC3M cells upon depletion of HNRNPA2B1 using a single siRNA duplex (si1, 20nM for 72 hours). **(F)** Log_2_ fold change gene expression values for differentially expressed *“Protein processing In endoplasmic reticulum”* genes upon HNRNPA2B1 depletion in PC3M cells (p<0.05 and absolute log_2_ fold change > 0.5 or < −0.5). P-values for each gene adjusted using the Benjamini and Hochberg method are represented by the bar colour – see key.

Given the previously established roles for *HNRNPA2B1* in the hypoxic response (Ho et al 2020, Yao et al 2013) and stress granule formation (Wolozin & Ivanov 2019), we wished to determine the most significant cellular stress pathways regulated by HNRNPA2B1 in PC. Firstly, we performed Gene Set Enrichment Class Analysis (GSECA) (Lauria et al 2020) on RNA-seq datasets from primary (n=491) (Hoadley et al 2018) and metastatic PC (CRPC) (n=208) (Abida et al 2019). We compared KEGG stress pathway representation in tumours with high *HNRNPA2B1* expression compared with low expression. In primary PC, we found that the top stress pathways associated with high expression of *HNRNPA2B1* included the *“Proteasome”* and *“HIF1 signaling pathway”* (Fig. 1C). In metastatic PC, top pathways associated with high expression of *HNRNPA2B1* included *“Protein processing in endoplasmic reticulum”, “Autophagy”*, and diseases with a misfolded protein component (Fig. 1D).

To validate these findings, we performed RNA-Seq of PC3M cells treated with either with a single siRNA duplex targeting *HNRNPA2B1* or a non-targeting control We observed a statistically-significant reduction in *HNRNPA2B1* gene expression following siRNA treatment as compared with the control (Log_2_ fold change = −4.05 adjusted p-value<0.001, Supplementary Table 5). Subsequently, we performed gene set enrichment analysis (GSEA) using all KEGG pathways to identify top biological processes enriched upon *HNRNPA2B1* depletion. Consistent with the association of HNRNPA2B1 with cellular stress pathways in PC patients, the KEGG stress pathway *“Protein processing in endoplasmic reticulum”* was the most significantly enriched pathway (Fig. 1E). Within this pathway, *HNRNPA2B1* depletion led to down-regulated expression of *PERK, ATF6* and *IRE1*, which encode for the three master ER-stress sensors mediating three key signaling branches of the UPR (Luo & Lee 2013) (Fig. 1F).

Taken together, these data in PC patients and cell lines indicates that HNRNPA2B1 regulates cellular stress pathways, with the most significant pathway being *“Protein processing in the endoplasmic reticulum”* in PC cells incorporating UPR genes.

### HNRNPA2B1 affects processing of IRE1 target mRNAs

To shed light on a putative mechanism of HNRNPA2B1-mediated UPR gene expression, we focussed on the IRE1-XBP1 signalling branch, considering its association with PC disease recurrence (Sheng et al 2019). XBP1 transcriptional activation requires non-canonical cytoplasmic splicing of XBP1u mRNA to produce the transcriptionally active XBP1s via removal of a variable 26 nucleotide sequence in exon 4 by IRE1 nuclease activity (Calfon et al 2002, Uemura et al 2009) (Fig. 2A). We hypothesised that HNRNPA2B1 may regulate UPR genes via XBP1 splicing. To test this, we used established RT-PCR based splicing assays (Savic et al 2014) to measure the percentage expression of activated XBP1s compared with XBP1u (Fig. 2A). Following treatment of PC3M cells with the UPR activator Thapsigargin (da Silva et al 2020); we observed a statistically significant increase in XBP1s splicing, compared to controls (Fig. 2B). Conversely, following HNRNPA2B1 protein depletion in PC3M cells using two independent siRNA duplexes (Fig. 2C); we observed a statistically significant decrease in XBP1s splicing compared with controls (Fig. 2D). These data demonstrate that HNRNPA2B1 promotes the non-conventional splicing of XBP1u to XBP1s.

**Figure 2.**
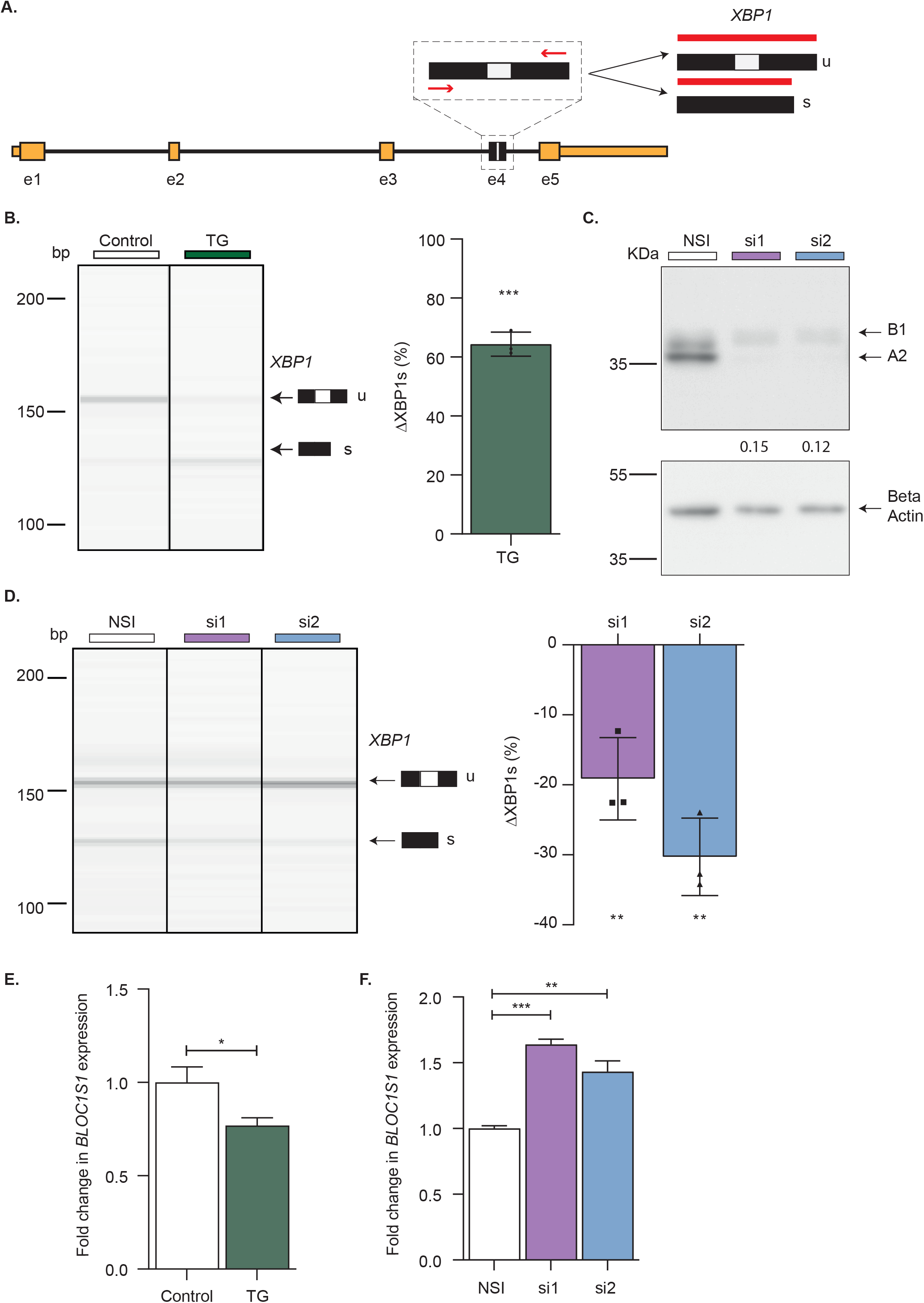
HNRNPA2B1 regulates processing of IRE1 target mRNAs. **(A)** Schematic of XBP1 gene. Exons 1-3 and 5 are indicated by yellow boxes and the non-canonically spliced exon 4 by a black box. XBP1u contains a variable 26-nucleotide region in exon 4 indicated by a white box, the exclusion of which generates the transcriptionally active XBP1s isoform. Red arrows represent RT-PCR primers used to amplify XBP1u and XBP1s products. **(B, left panel)** PC3M cells were treated with 250 nM Thapsigargin (TG), or vehicle (Control) DMSO for 24 hours and total RNA analysed using XBP1 splicing assays. Representative capillary gel electrophoretogram (QIAxcel) shows two bands representing transcripts with (XBP1u) or without (XBP1s) the exon 4 variable 26-nucleotide region inclusion**. (B, right panel)** Electrophoretograms were quantified to determine the percentage change in XBP1s product expression (ΔXBP1s). **(C)** PC3M cells were depleted of HNRNPA2B1 expression using two different siRNA duplexes (si1 and si2, 20nM for 72 hours) or non-silencing control (Nsi). Western blot shows HNRNPA2 (major isoform) and B1 (minor isoform) protein expression compared to Beta Actin loading control. The numbers below the HNRNPA2B1 blot indicate the relative reduction in total HNRNPA2B1 protein expression following siRNA depletion compared to Nsi control. **(D, left panel)** Total RNA was analysed using XBP1 splicing assays and representative capillary gel electrophoretogram show two bands representing XBP1u and XBP1s transcripts. **(D, right panel)** Electrophoretograms were quantified to determine the percentage change in XBP1s product expression (ΔXBP1s). **(E)** Relative change in *BLOC1S1* expression to DMSO control measured by qRT-PCR in PC3M cells treated with vehicle (Control) DMSO or Thapsigargin (TG) 250nM for 24 hours. **(F)** Relative change in *BLOC1S1* expression to Nsi measured by qRT-PCR in PC3M cells depleted of HNRNPA2B1 expression using two different single siRNA duplexes (si1 and si2, 20nM for 72 hours). At least three biological replicates were used, and Two-tailed T-test was used to compare treatment groups. * = p<0.05, ** = p<0.01, *** = p<0.001

IRE1 also degrades several mRNAs, including the BLOC1S1 mRNA, which encodes a regulator of lysosomal function, as part of the regulated IRE1-dependent decay (RIDD) pathway during ER stress (Chalmers et al 2019, Lhomond et al 2018). We wished to determine whether HNRNPA2B1 could also affect the RIDD pathway by exploring its impact on *BLOC1S1* expression. Following treatment of cells with the UPR activator Thapsigargin, we observed a statistically significant reduction in *BLOC1S* expression (Fig. 2E). Concordant with the impact of HNRNPA2B1 on XBP1 splicing, we observed a statistically-significant increase in *BLOC1S1* expression upon HNRNPA2B1 depletion (Fig. 2F). These data indicate that HNRNPA2B1 may affect multiple IRE1-dependent gene regulatory functions in PC cells.

### HNRNPA2B1-IRE1-XBP1 co-regulated genes represent a prognostic biomarker signature in primary PC and reveal a potential therapeutic target

Since high *HNRNPA2B1* expression is associated with poor PC patient prognosis (Fig. 1B), we hypothesised that this phenomenon may be mediated, in part, by HNRNPA2B1-dependent IRE1-XBP1-related gene expression. To test this, we utilised previously published RNA-Seq data from PC cells depleted of XBP1 or treated with the IRE1 inhibitor MKC8866 (Sheng et al 2019). To identify protein-coding genes co-regulated by XBP1, IRE1, and HNRNPA2B1, we overlapped lists of differentially expressed protein-coding genes in the three datasets (Fig. 3A). We identified a total of 20 HNRNPA2B1-IRE1-XBP1 co-regulated protein-coding genes.

**Figure 3.**
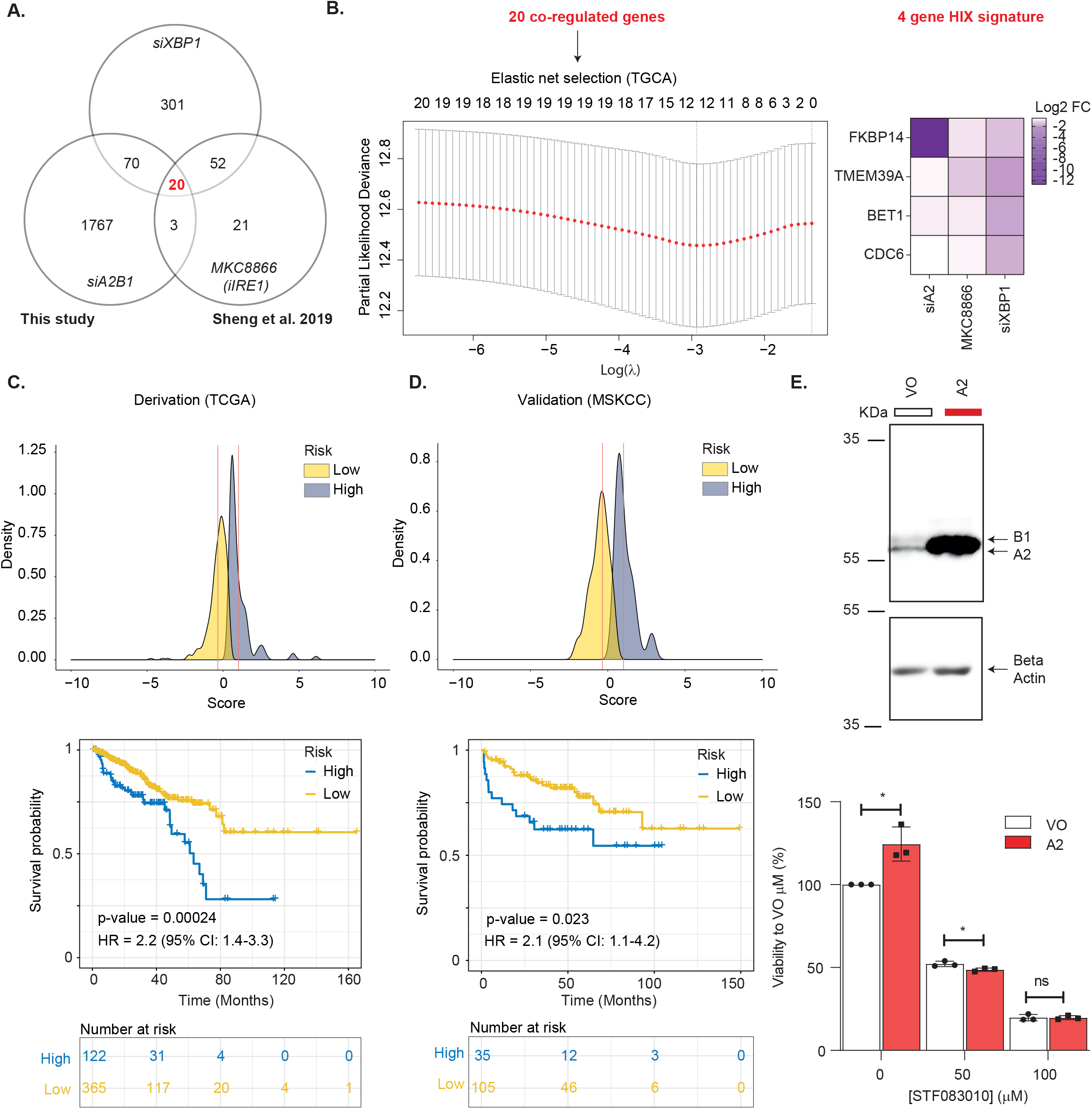
HNRNPA2B1-IRE1-XBP1 co-regulated genes represent a prognostic biomarker signature in primary PC and reveal a potential therapeutic target. **(A)** Venn diagram of protein-coding genes differentially-expressed and co-regulated by XBP1, IRE1 and HNRNPA2B1 with Log_2_ fold change <-0.5 and p<0.05 in RNA-Seq datasets from LNCaP cells treated with siRNA to XBP1 or IRE1 inhibitor MKC8866 (Sheng et al 2019) or PC3M cells treated with siRNA to HNRNPA2B1. **(B)** Derivation of prognostic biomarker panel by elastic net selection of 20 HNRNPA2B1, IRE1, and XBP1 co-regulated protein-coding genes in the TCGA cohort to generate a single four gene panel as the best predictors of disease relapse. **(B, left panel)** Cross-validation curve (red dots) with standard deviation. Left vertical dashed line is the value of λ that gives minimum mean cross-validated error (lambda.min), right vertical dashed line is the value of λ that gives the most regularized model such that the cross-validated error is within one standard error of the minimum (lambda.1se). **(B, right panel)** Heat map displaying the log_2_ fold change expression of the four HIX signature genes following treatment of LNCaPs with the IRE1 inhibitor MKC886, or XBP1 or HNRNPA2B1 depletion in LNCaP and PC3M cells respectively. **(C-D, top panels)** Distribution plot of risk scores for **(C)** derivation (TCGA) and **(D)** validation (MSKCC) cohorts. Vertical red lines represent mean of low and high percentile risk scores. **(C-D, bottom panels)** Kaplan-Meier plots of disease-free survival probabilities for patients from **(C)** derivation (TCGA) and **(D)** validation (MSKCC) datasets stratified by risk groups. The number of patients at risk for each group are presented in the table below each X-axis time point. Univariable Cox PH-dervied hazard ratios with 95% confidence intervals (CI) and two-tailed log-rank test p-values are shown. **(E)** PC3M cells were transfected with 3μg plasmid DNA (72 hours) encoding HNRNPA2 or vector only (VO) control. **(E, top panel)** Western blot shows HNRNPA2 protein expression compared to Beta Actin loading control. **(E, bottom panel)** PC3M cell viability was measured by MTT assay following transfection with 300 ng of plasmid DNA vector encoding HNRNPA2 or VO control. Cells were simultaneously treated with either 50 or 100 μM STF083010 or vehicle control (DMSO). Three biological replicates were used, and Two-tailed T-test was used to compare treatment groups. * = p<0.05

To determine if these 20 genes, or a subset thereof, were associated with disease recurrence, we performed elastic net regression using expression values of these genes in the TCGA cohort and time-to-event data (Fig. 3B). We applied elastic net selection (Fig. 3B, left panel, Supplementary Table 8) at lambda with the least mean cross-validation error and coefficient >0.00025 or <-0.00025. We identified four HNRNPA2B1-IRE1-XBP1 (HIX)-regulated genes *(FKBP14, TMEM39A, BET1*, and *CDC6)* as the best predictors of disease relapse (Fig 3B, right panel). Using multivariable Cox regression coefficient-derived patient risk scores for the four genes (see Materials and Methods), we stratified TCGA patients into two risk groups (low risk = <1st-3rd quartile, high risk = >3^rd^ quartile) (Fig. 3C, top panel). The high risk group was significantly more likely to relapse compared with the low risk group (Fig. 3C, bottom panel).

To validate these findings, we applied risk score calculations to an independent microarray-derived dataset (MSKCC) (Fig. 3D, top panel). Consistently, the high risk group was significantly more likely to relapse compared with the low risk group (Fig. 3D, bottom panel). Taken together, these data indicate that IRE1-XBP1-mediated gene activation may underlie the recurrent disease phenotype associated with HNRNPA2B1.

To determine whether the IRE-XBP1 signalling branch of the UPR might represent a potential therapeutic target for HNRNPA2B1 over-expressing PC, we firstly transiently ectopically expressed HNRNPA2, the predominant protein isoform encoded by *HNRNPA2B1* (Fig. 2C) in PC3M cells (Fig. 3E, top panel). Consistent with previously published data (Stockley et al 2014), we observed a statistically significant increase in cell growth following ectopic HNRNPA2 expression compared with controls (Fig. 3E, bottom panel). Subsequently, we treated HNRNPA2 overexpressing cells or controls with the IRE1 inhibitor STF083010 (Dong et al 2021). Following STF083010 treatment at 50 and 100μM doses, the effect of ectopic HNRNPA2B1 expression on cell growth was attenuated (Fig. 3E, bottom panel). These data suggest that IRE1 may be a potential therapeutic target in HNRNPA2B1 overexpressing PC tumours.

## Discussion

In this study, we reveal that high expression of HNRNPA2B1 in primary PC is associated with early disease recurrence. Our data indicate that this effect may mediated by HNRNPA2B1-controlled unfolded protein response (UPR) pathway-related genes via the major ER stress sensor IRE1. We show that HNRNPA2B1 controls IRE1-dependent XBP1 splicing and a subset of IRE1-XBP1 co-regulated genes classifies a subgroup of PC patients at high risk of disease relapse. Finally, we reveal that treatment with an IRE1 inhibitor attenuates HNRNPA2-driven PC cell growth, highlighting a novel line of therapy.

HNRNPA2B1 is known to play an important role in the formation of stress granules (Jiang et al 2021), and hypoxic adaptation (Ho et al 2020, Yao et al 2013). Here, we identify a link between *HNRNPA2B1* expression and several stress response pathways in primary and metastatic PC patients. In primary PC, we find that *HNRNPA2B1* largely is associated with metabolic stress pathways, whereas in metastatic PC it is associated with proteostasis stress such as *“Protein processing in endoplasmic reticulum”*. In tumourigenesis, sustained metabolic stresses, such as those caused by hypoxia, can disrupt proteostasis, induce ER stress, and activate the UPR (Ottens et al 2021). Hence, the association of *HNRNPA2B1* with *“Protein processing in endoplasmic reticulum”* in late-stage metastatic PC may be as a result of disrupted proteostasis acquired early in the disease course in a subset of patients with aggressive primary tumours over-expressing HNRNPA2B1.

Next, we reveal that HNRNPA2B1 regulates UPR gene expression including the master ER-stress sensor *IRE1*. Specifically, our findings implicates HNRNPA2B1 in IRE1-dependent processes of XBP1 splicing and RIDD activation. These two processes are mechanistically distinct, requiring dimerization or oligomerization of a phosphorylated version of the ribonuclease IRE1, respectively (Coelho & Domingos 2014). Given that depletion of HNRNPA2B1 increased expression of both *XBP1u* and *BLOC1S1*, we might speculate that HNRNPA2B1 may act downstream of IRE1 to regulate these mutually-exclusive events. Based on its known role in mRNA processing (Fahling et al 2006, Stockley et al 2014), it is possible that HNRNPA2B1 is either stabilises and/or facilitates transport of XBP1 and BLOC1S1 mRNAs to IRE1 at the ER membrane.

To identify a HNRNPA2B1-IRE1-XBP1-controlled prognostic biomarker signature (HIX) in PC patients, we initially used previously published RNA-seq datasets from PC cells treated with either the IRE1 inhibitor MKC8866 or depleted of XBP1 expression (Sheng et al 2019). Interestingly, both XBP1 siRNA and IRE1 inhibition regulate MYC protein expression and induce expression of several MYC target genes (Sheng et al 2019). Since MYC promotes the transcription of *HNRNPA2B1* (David et al 2010), we might speculate that HNRNPA2B1 is a component of the MYC-driven UPR activation.

XBP1 underpins several cancer hallmarks: XBP1 increases the key fatty acid metabolic enzyme SCD1 expression in MYC-driven cancers (Xie et al 2018). XBP1-mediated transcription of SNAI1, SNAI2, ZEB2, and TCF3 can mediate epithelial to mesenchymal transition and invasion (Cuevas et al 2017). By formation of a co-transcriptional complex with HIF1, XBP1 can controls angiogenesis (Chen et al 2014). Moreover, inhibitors of the IRE1-XBP1 pathway reduce tumour growth and sensitize cells to chemotherapy in pre-clinical models (Logue et al 2018, Sheng et al 2019). Here, we show that the known impact of XBP1 on PC cell growth and disease recurrence (Sheng et al 2019) is influenced by HNRNPA2B1.

Our study has several limitations: Although the novel link between HNRNPA2B1 and UPR was identified in metastatic PC, the HNRNPA2B1-IRE1-XBP1-controlled prognostic biomarker signature (HIX) was only validated in primary PC patients and based on mRNA expression. HNRNPA2B1 regulated *PERK* and *ATF6* as well as *IRE1* expression (Fig. 1F), however our validations focussed exclusively on IRE1-XBP1. Hence, we do not know the impact of HNRNPA2B1 on other UPR pathway branches. The precise molecular mechanisms underlying the HNRNPA2B1-mediated regulation of IRE1 and XBP1 remains unclear and warrants further investigation. Future studies using multiple UPR inhibitors in pre-clinical cancer models are required to determine whether targeting one or more UPR branches has therapeutic efficacy for HNRNPA2B1-overexpressing PC patients.

## Materials and Methods

### Transcriptomic datasets

Clinical RNA sequencing (RNA-Seq) and microarray data were obtained from cBioPortal (Cerami et al 2012, Gao et al 2013, Sanchez-Vega et al 2018). For primary PC (The Cancer Genome Atlas; TCGA, n=491 samples; Memorial Sloan Kettering Cancer Centre; MSKCC, n=179 samples), from Sanchez-Vega *et al*. (Sanchez-Vega et al 2018) for adjacent benign prostate (TCGA, n=52), and cBioPortal (Cerami et al 2012, Gao et al 2013) for metastatic PC (Stand Up to Cancer; SU2C, n=208 samples). Gene expression values were reported for TCGA as RNA-Seq by Expectation-Maximization (RSEM), for SU2C as Fragments per Kilobase of exon Per Million mapped fragments (FPKM) cohorts, or for MSKCC as log_2_ whole transcript mRNA expression. For comparison of normal (TCGA, n=52) and primary PC tissue (TCGA, n=497) RNA-Seq data were obtained from the Broad Institute Genome Data Analysis Center (GDAC) Firehose database (doi:10.7908/C11G0KM9) (Supplementary Table 1). Cell line RNA-Seq data for LNCaP cells treated with siRNA to XBP1 or and IRE1 inhibitor (MKC8866) were obtained from Sheng *et al*. (Sheng et al 2019) and gene expression values reported as Log_2_ Fold Change and adjusted p-value.

### Survival analysis

Patient samples were stratified into two groups by mRNA expression as follows: low = <1^st^-3^rd^ quartile and high = >3^rd^ quartile (Supplementary table 1). Kaplan-Meier plots were generated using time to event data (event = disease recurrence) from patient cohorts (TCGA; 487 out of 491 patients) using the *survfit* function of the *survminer* package in R V.4.1.1 and plotted using *ggsurvplot*. Univariable analyses were performed using the *coxph* function of *survminer*.

### Gene set enrichment class analysis (GSECA)

A list of 35 gene sets representing stress associated pathways was obtained from the Kyoto Encyclopaedia of Genes and Genomes (KEGG) pathway database https://www.genome.jp/kegg/pathway.html(Supplementary table 2). Patient samples were stratified into two groups by mRNA expression as follows: low = <1^st^-3^rd^ quartile and high = >3^rd^ quartile. GSECA was performed in R V.4.1.1 as previously described on stratified samples (Lauria et al 2020) considering the 35 KEGG stress associated pathway gene sets. An independent Monte Carlo simulation (1,000 iterations) was performed to determine the success rate (SR) of the association between the two cohorts. (Lauria et al 2020). Gene sets with GSECA association score (GAS) ≤0.05, adjusted p-value≤0.05 and success rate (SR)≥0.7 were considered as significant (Supplementary Table 2).

### Cell lines, transfections, and drug treatments

The PC3M cell line was generated as previously described (Pettaway et al 1996) and Short Tandem Repeat (STR) profiling (DDC Medical) used to confirm identity. Cells were maintained at sub-confluency, in RPMI-1640 medium (21875-034, Gibco) containing 2□mM L-glutamine, supplemented with 10% foetal calf serum (FCS) (Gibco), 100 units/ml penicillin and 100 μg/ml streptomycin (15140-122, Gibco), and incubated at 37°C, 5% CO_2_ in a humidified incubator. Cells were regularly screened for contamination with mycoplasma. DNA and siRNA transfections were performed as detailed in the figure legends using ViaFect (E4981, Promega) and RNAiMax (13778-075, Thermo Fisher Scientific), respectively, according to the manufacturers’ instructions (Supplementary Table 3). Cells were treated with IRE1 inhibitor (STF083010), at concentrations indicated in the figure legends, or vehicle control (DMSO).

### Antibodies, plasmids, and oligonucleotides

The plasmid pCAGPM-HA-hnRNPA2 (Katoh et al 2011) was a gift from Dr Y. Matsuura (Osaka University, Japan), and pcDNA3.1-HA was a gift from Professor T. Sharp (Queen Mary University of London, UK). The following antibodies were used: anti-HNRNPA2B1 (ab31645, Abcam), anti-actin (A1978, Sigma), anti-mouse IgG HRP-linked (P044701-2, Dako), and anti-rabbit IgG HRP-linked (P044801-2, Dako). The IRE1 inhibitor STF083010 was purchased from Merck Life Science, UK (SML0409). Sequences used to generate siRNA duplexes are as previously described (Stockley et al 2014) or commercially-designed (ON-TARGETPlus, Dharmacon Horizon Discovery) (Supplementary Table 3). Primers for PCR were designed using the National Center for Biotechnology Information (NCBI) Primer-BLAST tool (https://www.ncbi.nlm.nih.gov/tools/primer-blast) with the Ensembl (http://www.ensembl.org) Transcript ID for the principal mRNA isoform and synthesised by Integrated DNA Technologies (Supplementary Table 3). Primers for *in vitro* splicing analysis are as previously published (Savic et al 2014) (Supplementary Table 3).

### SDS-PAGE and Western blotting

Whole cell lysis was performed in RIPA (Radio-Immunoprecipitation Assay) buffer for 30 minutes at 4°C. Protein concentration was determined by Bicinchoninic acid (BCA) assay (10678484, Thermo Fisher Scientific) and samples adjusted to the same total protein concentration. Proteins were separated by size by SDS-Polyacrylamide Gel Electrophoresis (SDS–PAGE) on 10% w/v gels and electroblotted onto a Polyvinylidene fluoride or polyvinylidene difluoride (PVDF) membrane (3010040001, Sigma). Luminata Crescendo Western HRP substrate (10776189, Thermo Fisher Scientific) was used for signal detection, and protein bands were visualised on a Chemidoc system (Amersham Imager 600, Amersham). Antibody concentrations were as follows: anti-HNRNPA2B1 (1:1 000), anti-actin (1:100 000); HRP-linked secondaries (1:5 000). Where indicated, densitometry assessments of protein bands were performed using Image Studio Lite v.5.2 (LI-COR), and signal intensities used to calculate relative normalised fold-change in protein expression (Supplementary Table 4).

### RNA-Seq and gene set enrichment analysis

Total RNA was extracted from cells using the QIAgen RNeasy mini kit (74004, QIAgen), and treated with DNAse I (AMPD1, Sigma) to exclude genomic contamination. Libraries were generated using the TruSeq RNA Library Prep Kit v2 (RS-122-2001, Illumina) and 75bp paired end sequencing performed to 30M read depth using the NextSeq 500 (Illumina). Reads were aligned to the genome (hg38) using STAR v2.7.3a in dual pass mode. Transcripts were assembled and quantified in Transcripts Per Million (TPM) using Stringtie v2.1.1 (Pertea et al 2015). Read normalisation and differential gene expression analysis was performed using DESeq2 v1.34.0 in R V.4.1.1 (Supplementary Table 5). Enrichment of KEGG pathways amongst differentially-expressed genes with log_2_ fold change of <-0.5 or >0.5 at p<0.05 significance was performed in R V.4.1.1 using the *enrichKEGG* function of the *clusterProfiler* package (Wu et al 2021) in R and plotted with *dotplot* (Supplementary Table 5). Raw data have been deposited at Gene Expression Omnibus (http://www.ncbi.nlm.nih.gov/geo) under accession number GSE198261, and all details are Minimum Information About a Microarray Experiment (MIAME) compliant.

### Quantitative Reverse Transcription PCR

Total RNA was extracted from cells using TRI Reagent Solution (M9738, Thermo Fisher Scientific), and reverse transcribed to cDNA using the High Capacity cDNA Reverse Transcription Kit (4368814, Applied Biosystems). cDNA (20ng per condition) was combined with forward and reverse primers (Supplementary Table 3) and the Luna Universal qPCR Master Mix master mix (M3003, NEB) containing SYBR green and ROX passive dye to a final 10ul reaction volume. Binding of SYBR green to DNA was analysed in a QuantStudio 5 Real-Time PCR system (Thermo Fisher Scientific). Reaction conditions were as follows: Initial denaturation at 95°C for 10 minutes, 40 cycles of denaturation for 15 seconds at 95°C, plus annealing, extension, and signal capture at 60°C for 1 minute. The 2-ΔΔCT method was used to determine relative gene expression using the geometric mean expression of two validated endogenous control genes *(ACTB* and *B2M)* (Supplementary Table 6).

### XBP1 Splicing Assays

Total RNA was extracted from cells using TRI Reagent Solution (M9738, Thermo Fisher Scientific), and reverse transcribed to cDNA using the High Capacity cDNA Reverse Transcription Kit (4368814, Applied Biosystems). cDNAs (20ng per condition) were combined with primers flanking the variable exonic region of XBP1 (Savic et al 2014) (Supplementary Table 3), dNTPs and Taq Polymerase (NEB, M0273) in standard reaction buffer to a final 10ul reaction volume. Reactions were performed in a ProFlex thermocycler (Applied Biosystems) as follows: Initial denaturation at 95°C for 30 seconds, 30 cycles of denaturation for 15 seconds at 95°C, plus annealing at 52°C for 30 seconds, and extension at 68°C for 1 minute; followed by a final extension at 68°C for 5 minutes. PCR products were resolved, detected and quantified by capillary gel electrophoresis (QIAxcel, QIAgen) (Supplementary Table 7).

### Derivation and validation of a prognostic biomarker panel

To identify the combination of genes which are the strongest predictors of PC recurrence, the *glmnet* package (Friedman et al 2010) in R V.4.1.1 was used to fit gene expression to time-to-event data in the TCGA (derivation) cohort using cox regression with an α = 0.2using a coefficient cut off of >0.00025 or <-0.00025 at λ minimum. To obtain coefficients representing the relative contributions of the selected genes to the prognostic value of the signature, multivariable analysis was performed using time-to event data and grouped expression of each of the four signature genes (low = <1^st^-3^rd^ quartile and high = >3^rd^ quartile), by the *coxph* function of *survminer* package. Coefficients for each gene were obtained from the high expression group (Supplementary table 8). Next, a risk score (i) for each patient was derived from the coefficients of the multivariable Cox PH model as follows: 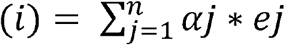, where αj is the scaled j gene expression value with ej coefficient in the derivation multivariable model (Royston & Altman 2013). Risk group cut-offs were defined based upon quartiles of gene signature score in TCGA data (low = <1^st^-3^rd^ quartile, high = >3^rd^ quartile).

Kaplan-Meier plots were generated using time to event data (event = disease recurrence) from patient cohorts using the *survfit* function of the *survminer* package in R V.4.1.1 and plotted using *ggsurvplot* (Supplementary table 8). Univariable analyses were performed using the *coxph* function of *survminer* to compare patients with low and high risk scores. To validate the model, risk scores calculated using the coefficients obtained from the derivation cohort were applied to scaled gene expression values from the validation cohort, and Kaplan Meier plots generated stratified by risk scores (low = <1^st^-3^rd^ quartile, high = >3^rd^ quartile).

### Cell growth assay

Cells (n = 2000) were seeded into each well of a 96-well plate and grown to ~20–30% confluence prior to transfection with DNA as indicated in the figure legends. After 72 hours, (3-(4,5-Dimethylthiazol-2-yl)-2,5-Diphenyltetrazolium Bromide) (MTT) (M6494, Thermo Fisher Scientific) was added to each well to a final concentration of 0.67 mg/ml and incubated at 37oC, 5% CO2 in a humidified incubator for 2 h. MTT reagent was then removed, and 100μl dimethyl sulfoxide (DMSO) (10213810, Thermo Fisher Scientific) added to each well, and the plate was agitated at room temperature for 15 minutes. Absorbance was measured at 560nm and 630nm (SpectraMax Plus384 Absorbance Microplate Reader, Molecular Devices), and normalized by subtracting the 630nm value from the 560nm value. Percentage viability (%) was calculated as: the treatment absorbance divided by the DMSO control absorbance. All data were normalized to a vector only control (Supplementary Table 9).

## Data Availability

RNA-Seq data from this publication have been deposited to Gene Expression Omnibus and assigned the identifier accession number GSE198261.

## Acknowledgements

We would like to thank Y. Matsuura (RIMD, Japan) and T. Sharp (QMUL, UK) for providing plasmid DNA vectors used in this study. We are grateful to the P. Herzyk, J. Galbraith, G. Hamilton, and M. Mudaliar (University of Glasgow Polyomics, UK) as well as A. Hedley and G. Kalna (CR-UK Beatson Institute, UK) for assistance with RNA-seq and bioinformatics. We would also like to thank P. Grevitt (QMUL, UK) and P. Baptista-Ribeiro (QMUL, UK) for their critical appraisal of earlier versions of the manuscript. The research performed in this study was funded by the Royal College of Surgeons of England/Cancer Research UK Clinician Scientist Fellowship in Surgery (C19198/A15339 to PR), The Urology Foundation and John Black Charitable Foundation (to PR), Barts Charity (MGU0533 to PR) and Orchid Charity (to PR).

